# Quantitative Systems Pharmacology Model for TROP-2 Targeted Antibody-Drug Conjugate in Triple-Negative Breast Cancer

**DOI:** 10.64898/2026.09.20.752985

**Authors:** Sedat Dogru, Ashwatha Suresh, Edward P. Bowman, Anshu Marathe, Azher M. Hussain, Aleksander S. Popel

## Abstract

TROP2-targeted antibody-drug conjugates (ADCs) have demonstrated promising clinical activity in triple-negative breast cancer (TNBC) as monotherapies; however, therapeutic benefit varies among patients. Combination strategies pairing TROP2-targeted ADCs with immune checkpoint inhibitors are also being investigated. Elucidating the mechanistic drivers of ADC monotherapy variability and enabling the rational development of combination regimens require computational frameworks that integrate ADC pharmacology with tumor-immune interactions. A quantitative systems pharmacology (QSP) model is presented that incorporates an ADC module into our established immuno-oncology model for TNBC. The module captures ADC and payload pharmacokinetics and pharmacodynamics. TNBC heterogeneity is represented by two tumor cell clones with high and low TROP2 expression, informed by prior characterizations, and differential sensitivity to the ADC payload is incorporated as an intrinsic property of each clone. Although generalizable, the model was applied to the TROP2-targeted ADC sacituzumab govitecan (SG, TRODELVY). A virtual patient cohort was generated using Latin hypercube sampling and calibrated against objective response rate (ORR) data from SG’s Phase I/II TNBC basket trial. The model predicted an ORR of 33.2% consistent with ASCENT study (NCT02574455). Simulations suggest TROP2-mediated delivery contributes modestly to SG efficacy with tumor exposure driven largely by systemically released SN-38 payload being sufficient to induce cytotoxicity. Tumor heterogeneity emerged as a key determinant of response with ORR increasing as the fraction of payload-sensitive clones increased. Overall, this QSP framework for TROP2-targeted ADCs accounts for TNBC heterogeneity and is extendable to other ADCs and targets enabling interrogation of ADC mechanisms of action in conjunction with tumor-immune interactions.

## 1. Introduction

Antibody-drug conjugates (ADCs) are emerging as a promising therapeutic approach in oncology.^1^ They are composed of three main components: an antibody, a linker, and a payload which combine the specificity of monoclonal antibodies with the cytotoxic potential of small-molecule drugs.^1–3^ They have the potential of delivering cytotoxic small molecules to tumor microenvironment with minimal systemic exposure potentially decreasing or preventing adverse events caused by chemotherapies. There are currently 14 FDA-approved ADCs for different cancer indications.^1^

Triple-negative breast cancer (TNBC) is associated with a poor prognosis and high recurrence rates.^4,5^ Trophoblast cell surface antigen 2 (TROP2) is an attractive therapeutic target for ADC delivery in TNBC as it is highly expressed in cancer cells.^6–9^ Therefore, TROP2-targeted ADCs are emerging as a promising therapeutic approach in TNBC with one approved (sacituzumab govitecan, SG) and several other ADCs in late clinical trial stages such as Dato-DXd and ESG-401.^8,10,11^ Clinical outcomes of ADCs in TNBC patients remain highly variable with a limited understanding of the underlying reasons.^11–15^ While TROP2 is highly expressed in cancer cells, its expression levels exhibit heterogeneity among TNBC cells. TROP2 expression varies considerably across TNBC molecular subtypes with cells categorized in mesenchymal-like subtype typically exhibiting low TROP2 expression and basal-like subtype cells displaying higher levels.^5,16–18^ The expression difference may limit TROP2-targeted ADC efficacy in patients with mesenchymal subtype–dominant tumors or in tumor regions with mesenchymal-like characteristics. Notably, mesenchymal-like cell lines are also reported to be more resistant to DNA-damaging agents (DDAs), including topoisomerase I inhibitors such as SN-38 and DXd, compared to basal-like cell lines.^19–21^ These findings suggest a covariance of TROP2 expression and payload sensitivity where tumor regions with higher TROP2 expression may also have greater sensitivity to the delivered payload, and regions with lower TROP2 expression may have lower sensitivity. Overall, these findings suggest that heterogeneity in both TROP2 expression and payload sensitivity may play a role in the efficacy of TROP2-targeted ADCs. Therefore, understanding their contributions to overall efficacy is necessary to potentially improve therapeutic outcomes.

Quantitative systems pharmacology (QSP) models provide a mechanistic framework for *in silico* exploration of complex biological mechanisms and variability in therapeutic response.^22,23^ Multiple QSP models have been published in recent years focusing either on specific antibody-drug conjugates (ADCs) or on more general modeling frameworks.^24–26^ These models have supported ADC design and improved understanding of their mechanisms of action. A critical aspect of developing a QSP model for ADCs is the representation of tumor heterogeneity. The heterogeneity has multiple layers: (1) target antigen expression heterogeneity among cancer cells and (2) variability in cellular sensitivity to the payload. The first factor is important because it impacts the distribution of the ADC within the tumor.^27,28^ The second factor is important because, even when the payload is successfully delivered, its efficacy depends on the sensitivity of the cancer cells to the payload. In current QSP ADC models, tumor heterogeneity is typically represented by antigen expression heterogeneity (case 1); whereas, variability in payload sensitivity (case 2) is often overlooked despite its potential importance for ADC analysis due to bystander effects. In addition, existing models have generally been developed with a primary focus on ADCs without explicitly including immune-cancer interactions. Current trends indicate that combination therapies involving ADCs and immune checkpoint inhibitors are promising with an increasing number of clinical trials underway.^29^ Therefore, a comprehensive model that integrates both an ADC module and immune-cancer interactions is required.

Our aim was to develop a mechanistic QSP model that integrates a TROP2-targeted ADC module with immune-cancer interactions, while explicitly representing TNBC heterogeneity, to understand the mechanisms underlying ADC activity in TNBC and its potential synergy with the immune response. The previously developed QSP immuno-oncology model was extended by incorporating a TROP2-targeted ADC module for TNBC to achieve this aim.^23^ Importantly, this approach explicitly accounts for heterogeneous tumor populations at two levels by distinguishing antigen-high and antigen-low clones which affects ADC delivery and by incorporating their relative sensitivity to the payload which influences the cytotoxic efficacy of the payload after delivery. This approach provides a comprehensive framework for mechanistic exploration of ADC mechanisms of action, their interaction with the immune response, and the role of TNBC heterogeneity in determining ADC efficacy.

SG was used as a representative compound to parameterize, calibrate, and validate our QSP model in this study. It received accelerated FDA approval for TNBC in 2020 by achieving overall response rates of 31% and a disease control rate of 67%.^30,31^ It incorporates a humanized anti-TROP2 antibody (sacituzumab), a pH-sensitive linker (CL2A), and a topoisomerase I inhibitor payload (SN-38) with an average drug-to-antibody ratio (DAR) of 7.6.^32^ SG exhibits several distinctive features. First, it has a relatively short half-life (16 hours) compared with many FDA-approved ADCs (4–6 days).^32,33^ Second, its payload SN-38 is approximately an order of magnitude less potent than other commonly used ADC payloads such as MMAE and DM1.^32^ The lower potency allows administration of the highest dose (10 mg/kg) of any FDA-approved ADCs. Furthermore, its pH-sensitive linker can undergo spontaneous hydrolysis in circulation leading to systemic SN-38 release.^34^ The contribution of these distinctive features or their interplay to SG’s efficacy is not well understood. As an application example of the proposed framework, we parameterized the model for SG, calibrated the model using SG’s Phase I/II clinical trial data, and performed validation using data from SG’s Phase III ASCENT clinical trial.

## 2. Methods

The QSP model comprises four main compartments (central, peripheral, tumor-draining lymph node, and tumor) and includes 287 ordinary differential equations (ODEs), 42 algebraic equations (i.e., repeated assignment rules), and 308 parameters (Fig. 1). Model species, parameters, reactions, compartments, and algebraic equations are listed in the supplemental information (Table S1-S5, respectively) and are explained in detail in our previous studies.^23,35^ An ADC module was developed in the present study and incorporated into our previously published QSP model.

**Fig. 1.**
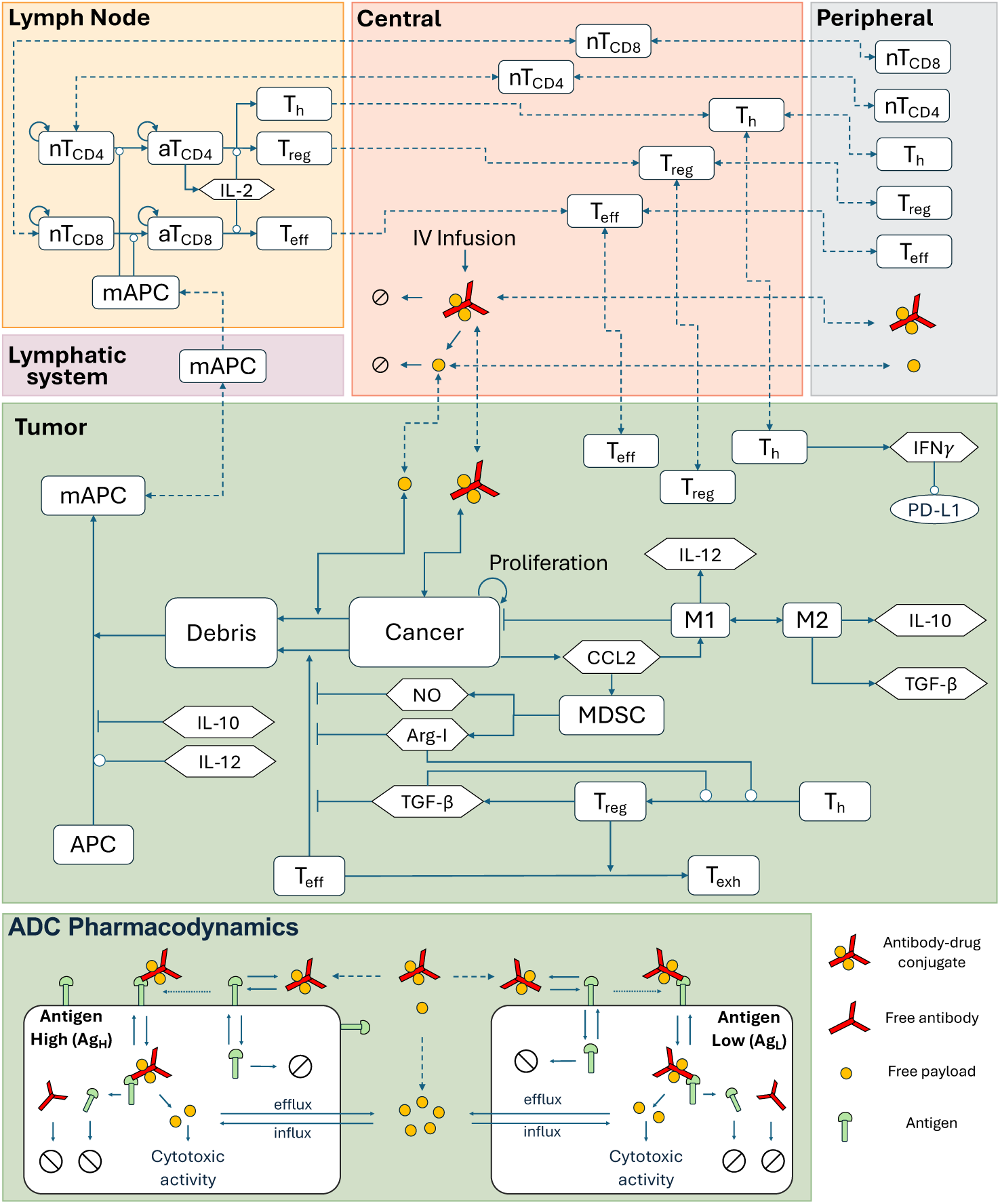
Schematic of the QSP model, which consists of four main compartments: central, peripheral, tumor, and tumor-draining lymph node. Each compartment is designed in a modular manner with submodules. The ADC module is incorporated as a submodule of the tumor compartment. The ADC is administered in the central compartment, where it can release its payload or transport into the peripheral or tumor compartments. Payload released in the central compartment can transport into the peripheral or tumor compartments. In the tumor compartment, the ADC can bind to its target antigen, become internalized, and release the payload intracellularly. The released payload can induce cytotoxicity or efflux into the tumor microenvironment. nT: naive T cell; aT: activated T cell; NO: nitric oxide; Arg-I: arginase I; Treg: regulatory T cell; Teff: effector T cell; Th: helper T cell; M1&M2: M1 and M2 like macrophage; APC: antigen-presenting cell; mAPC: mature antigen-presenting cell. White circle and T-bar denote stimulation and inhibition, respectively.

### 2.1. ADC Module

The ADC module accounts for ADC-TROP2 complex kinetics (binding, internalization, intracellular toxin release, and denaturation) and unconjugated toxin kinetics (cytotoxicity, influx, and efflux). SG was used as a representative TROP2-targeted ADC to test and calibrate the model. Parameterization of the model was performed using published SG in vitro and in vivo data. All parameters and their derivation are explained in the supplemental information (Fig. S1-S3). SG pharmacokinetics were adapted from an earlier study.^34^ Briefly, a two-compartment model was employed for SG and SN-38. SG is administered to the central compartment at 10 mg/kg on Day 1 and Day 8 of a 21-day cycle and SN-38 release follows first-order kinetics driven by SG concentration. An apparent clearance rate was used for SN-38 due to identifiability issues. The PK model parameters are provided in the supplemental information.

### 2.2. TNBC Heterogeneity

TNBC heterogeneity was represented by two distinct cell clones in the ADC module: high and low TROP2-expressing populations with different sensitivities to SN-38. We represented previously established TNBC subtype-specific TROP2 heterogeneity in which cell lines in the basal-like molecular subtype category are characterized by higher TROP2 expression levels compared to mesenchymal-like molecular subtypes (Fig. S4).^36^ Therefore, these subtypes were represented as antigen-high (Ag_H_) and antigen-low (Ag_L_) cell clones. Furthermore, several studies have reported that while basal-like subtype cells are sensitive to topoisomerase inhibitors such as SN-38, mesenchymal-like subtypes are resistant.^20,37^ To represent this in our model, we used reported potency (IC50) values for cell lines in both subtypes and implemented the median, lower threshold, and higher threshold values in the model.^38^ Additionally, cytotoxicity rates (E_max_) for both subtypes were approximated from in vivo xenograft tumor growth inhibition experiments (Fig. S3).

### 2.3. *In Silico* Clinical Trials

A virtual patient (VP) cohort was generated using Latin Hypercube Sampling of 34 parameters (Table S6) and calibrated against the response status outcomes of TNBC patients in SG’s Phase I/II basket trial (NCT01631552) using Bayesian optimization algorithm (bayesopt) in Matlab. Twenty-six parameters were adopted from our previous studies^23,39,40^ selected to represent interpatient variability and eight additional ADC-related parameters were introduced to capture potential sources of variability specific to SG. The complete list of parameters and their values is provided in the supplementary information (Table S6). Non-evaluable patients in the clinical trial were excluded and response status rates were calculated based on reported numbers of partial and complete response (PR/CR), stable disease (SD), and progressive disease (PD) patients. In the QSP model, the response status of each virtual patient was evaluated by the change in tumor volume based on RECIST 1.1.^41^ To calibrate our QSP model, we first tested the effect of the number of VPs on response status estimations via step by increase of the number of VPs (n= 100-2400) using bootstrap confidence intervals. No significant difference was observed beyond 400 VPs (Fig. S5A). Therefore, the calibration was performed with an initial cohort size of 400 VPs. Subsequently, we tested the calibrated parameter distribution for estimating Phase III clinical trial (ASCENT, NCT02574455) response status results using bootstrap confidence intervals (n=244) without further calibration of the model. Simulations were performed in MATLAB SimBiology Toolbox (MathWorks, Natick, MA) using the Sundials solver, with absolute and relative tolerances of 1e-9 and 1e-6, respectively.

### 2.4. Statistical Analysis

Latin Hypercube Sampling and Partial Rank Correlation Coefficient (LHS-PRCC) methods were employed for global sensitivity analysis to examine the impact of varied parameters on model observations.^42^ The Top-Down Correlation Coefficient (TDCC), which is based on Savage scores, has been proposed in the statistical literature for LHS-PRCC analysis to determine the number of VPs required to obtain statistical power for the global sensitivity analysis.^42^ Briefly, we used TDCC analysis and tested increasing VP cohort sizes from 100 up to 2400. The correlation coefficient between subsequent VP sizes plateaued around 1200 VPs, which we took as a reasonable point of stability (Fig. S5B). Therefore, the rest of the simulations were performed with 1200 VPs.

## 3. Results

### 3.1. Prediction of SG PK and Payload Distribution in Plasma and Cancer Tissue

The distribution of SG and its payload, SN-38, in the central and tumor compartments over time was predicted. SG concentrations reached maximum concentration of 2.2 × 10⁵ ng/mL and 4.9 × 10⁴ ng/mL in the central and tumor compartments, respectively, which is consistent with clinical trial observations (Fig. 2A).^34^ Systemically released SN-38 concentrations reached approximately 96.5 ng/mL and were able to transport into the tumor compartment. Free SN-38 concentrations in the tumor compartment reached 42.7 ng/mL and remained above 20 ng/mL for 24 hours (Fig. 2B).

**Fig. 2.**
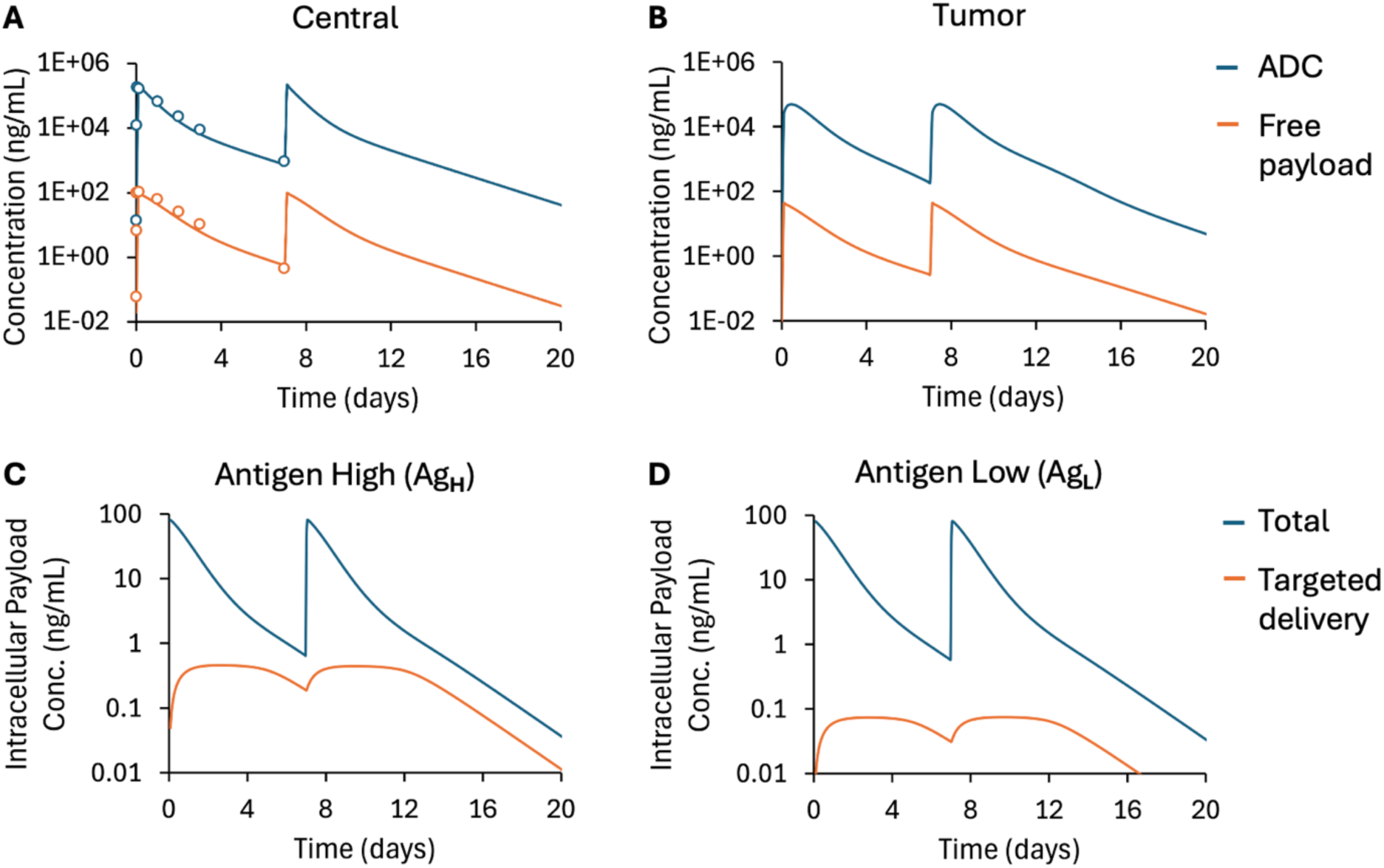
A two-compartment PK model was employed to predict the time-dependent concentrations of the ADC and free payload in the (A) central and (B) tumor compartments. SG at 10 mg/kg was administered on Days 1 and 8 of a 21-day cycle. Contribution of targeted ADC delivery to intracellular payload concentrations was quantified by isolating the payload delivered through SG-TROP2 complex internalization and subsequent intracellular payload release for both (C) Ag**_H_** and (D) Ag**_L_** cell clones. A small fraction of the intracellular payload originates from SG-TROP2 internalization, indicating that most of the intracellular payload is sourced from the systemically released portion, transported to the tumor compartment, and subsequently taken up by tumor cells.

Although systemically released SN-38 corresponds to approximately 4.5% of the total administered SN-38 after 24 hours with about 95.5% remaining conjugated,^43^ the free SN-38 levels in plasma may be sufficiently high to induce cytotoxic effects if transported into the tumor microenvironment. Therefore, we analyzed the contributions of both SN-38 sources to the total SN-38 delivered to tumor cells: payload delivered via internalization of the SG-TROP2 complex and subsequent intracellular release and free SN-38 in plasma permeating into the tumor microenvironment. We first isolated the payload delivered exclusively through SG-TROP2 complex internalization and subsequent intracellular release in both Ag_H_ and Ag_L_ cell clones by disabling plasma-to-tumor SN38 permeability in the model. The resulting SG-TROP2 mediated intracellular SN38 exposure corresponded to approximately 2.4% and 0.4% of the total cellular SN38 exposure observed in Ag_H_ and Ag_L_ clones, respectively, when plasma-to-tumor SN38 permeability was enabled (Fig. 2C-D). These results indicate that only a small fraction of total intracellular exposure arises from targeted delivery; whereas, most of the intracellular payload originates from the systemically released fraction that is transported to the tumor compartment and subsequently taken up by tumor cells. Consequently, while targeted delivery differs between Ag_H_ and Ag_L_ cell clones, the total intracellular exposure is similar in both clones, thereby reducing the impact of antigen-expression heterogeneity on treatment outcomes.

### 3.2. Tumor Growth Inhibition in heterogeneous TNBC

The effect of TNBC heterogeneity on tumor growth inhibition was analyzed by varying the initial fractions of Ag_H_ and Ag_L_ cell clones. The model suggests that TNBC heterogeneity strongly influences the tumor growth inhibition efficacy of SG. In the presence of only the Ag_H_ cell clone, cancer cells were completely eliminated within less than six months (Fig. 3A), and the tumor fully regressed (Fig. 3C). However, increasing the initial fraction of the Ag_L_ cell clone within the tumor reduced the efficacy of SG (Fig. 3B); even an initial Ag_L_ fraction of 25% led to tumor relapse and renewed growth after approximately three months (Fig. 3C).

**Fig. 3.**
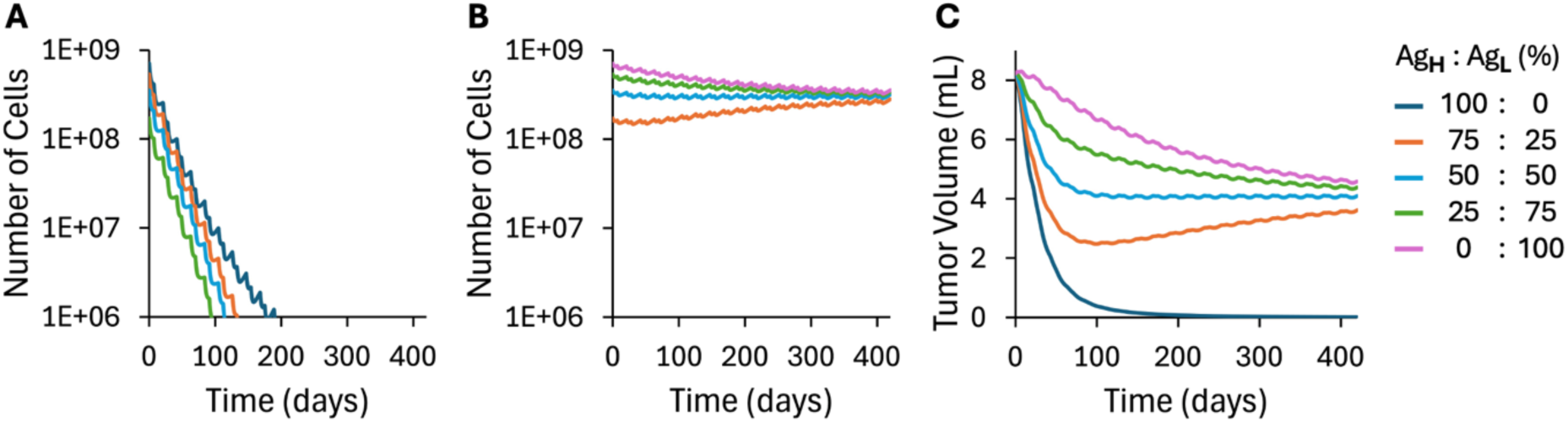
Tumor growth-inhibition efficacy of SG was evaluated across varying levels of TNBC heterogeneity, represented by the initial fractions of antigen-high (Ag**_H_**) and antigen-low (Ag**_L_**) cell clones. Changes in the number of (A) Ag**_H_** and (B) Ag**_L_** cells, as well as (C) the overall tumor volume over time, were estimated for different initial Ag**_H_**:Ag**_L_** fractions. Increased tumor growth inhibition with higher Ag_H_ fractions may suggest a TROP2-mediated increase in ADC delivery; however, as shown in Fig. 2C-D, TROP2-mediated delivery accounts for only a small fraction of the delivered payload. The increased tumor growth inhibition is instead driven by a higher fraction of payload-sensitive clones (Ag_H_), which correlates with TROP2 expression levels.

### 3.3. Virtual Patient Cohort Generation and In Silico Clinical Trials

The calibrated virtual patient cohort was used to simulate tumor-growth inhibition by SG in TNBC. Simulations were performed for 1200 VPs and those that did not reach the initial tumor diameter criteria (>1 cm) were excluded. Parameter sets of the remaining VPs were saved for subsequent simulations of therapy-dynamics and statistical analysis. Model predictions showed good agreement with clinical trial response status: PR/CR 34.0% vs 33.2%, SD 39.3% vs 38.5%, and PD 26.6% vs 28.3% for clinical trial versus simulation, respectively (Fig. 4A). Confidence intervals were calculated for 244 evaluable VPs.

**Fig. 4.**
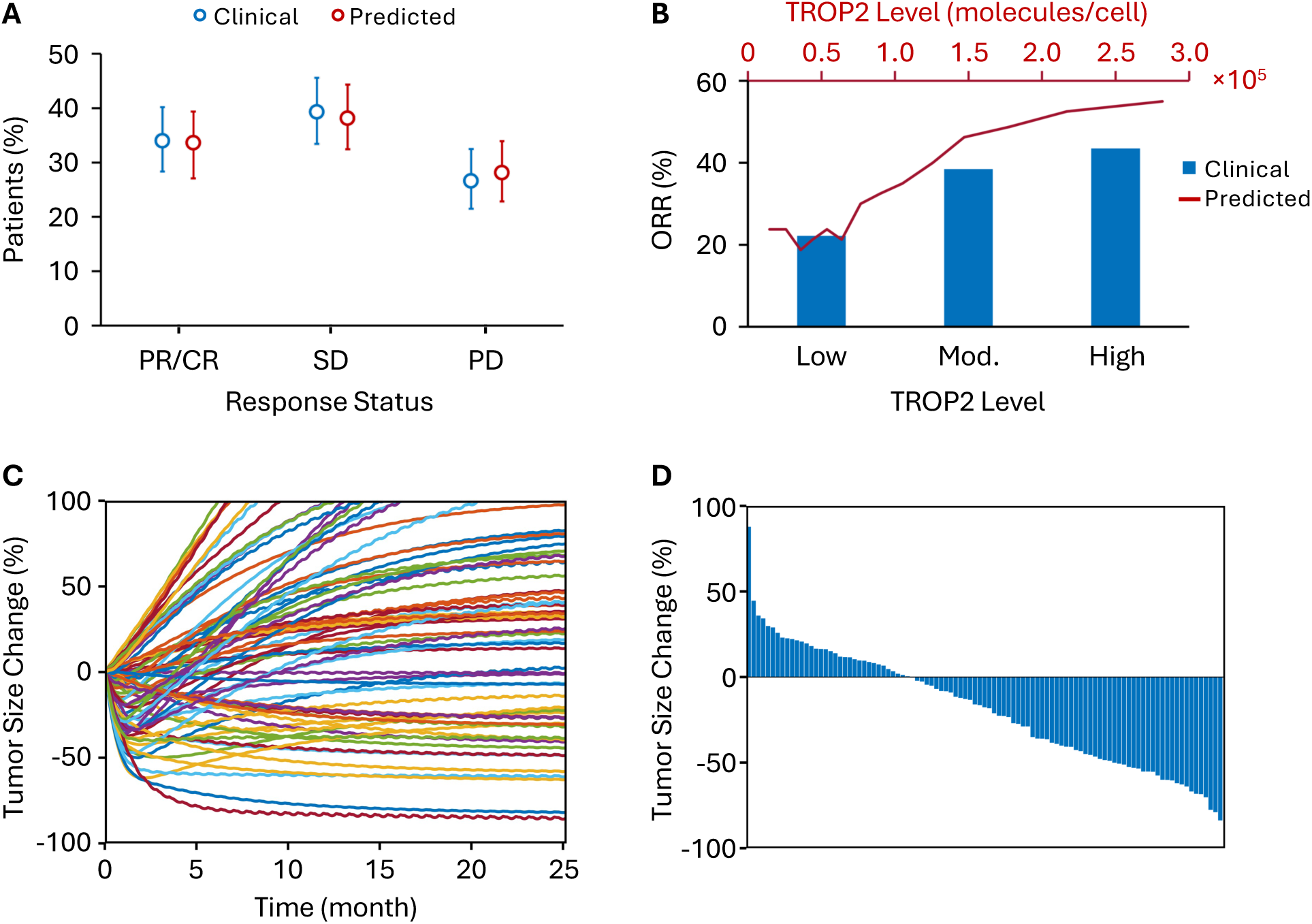
Comparison of clinical and model-predicted patient responses and tumor size dynamics. (A) Comparison of the percentage of patients in different response categories: Partial Response/Complete Response (PR/CR), Stable Disease (SD), and Progressive Disease (PD), between clinical observed data and model-predicted results. Error bars represent the 95% confidence interval. (B) Comparison of overall response rate (ORR, %) stratified by TROP2 expression levels (low, moderate [Mod.], and high) between clinical data ^44^ and model predictions. Although TROP2 levels appear to be associated with response, their contribution to payload delivery is limited, as shown in Fig. 2C–D. Therefore, the observed increase in ORR with TROP2 reflects the covariation between TROP2 expression and tumor sensitivity to SN38. (C) Spider plot showing individual virtual patient tumor size change (%) over time. Each line represents a single patient’s tumor size trajectory. (D) Waterfall plot showing the best overall tumor size change from baseline for each individual patient. Both (C) and (D) show results for 100 randomly selected virtual patients.

Analysis further revealed that ORR increased with TROP2 expression, rising from 23.8% in the lowest to 55.0% in the highest TROP2-expressing virtual patient group, consistent with trends observed in clinical biomarker analyses (Fig. 4B).^44^ Although TROP2 levels appeared to be associated with response, their contribution to payload delivery was limited, as shown in Fig. 2C-D, and thus they do not play a major role in driving efficacy. Instead, the increase in ORR with TROP2 reflects the covariation between TROP2 expression and tumor sensitivity to SN38. Time-dependent percentage changes in tumor size (spider plots) for 100 randomly selected VPs based on RECIST criteria are shown in Fig. 4C. A waterfall plot depicting changes from baseline in model-predicted tumor diameter for 100 randomly selected VPs is presented in Fig. 4D.

#### 3.3.1. Determinants of SG Therapeutic Efficacy

A global sensitivity analysis was performed to identify the key drivers of SG therapeutic efficacy within the QSP model. PRCCs were computed to assess the impact of model parameters on predicted tumor volume over time. The ranked sensitivities of the most influential parameters are presented in Fig. 5A. The analysis revealed that tumor volume was highly sensitive to parameters governing tumor vasculature dynamics and pharmacodynamics of the SN-38 payload throughout the simulation. Specifically, Ag_L_ SN-38 cytotoxicity rate constant (E_max_) and the tumor vasculature inhibition rate displayed the strongest negative correlations with tumor volume. Conversely, the tumor vasculature growth rate and Ag_L_ SN-38 potency (IC50) showed strong positive correlations with tumor volume throughout the simulation indicating that higher values of these parameters are associated with reduced efficacy. Similarly, the initial tumor diameter displayed a strong early positive correlation reflecting its direct contribution to baseline tumor burden.

**Fig. 5.**
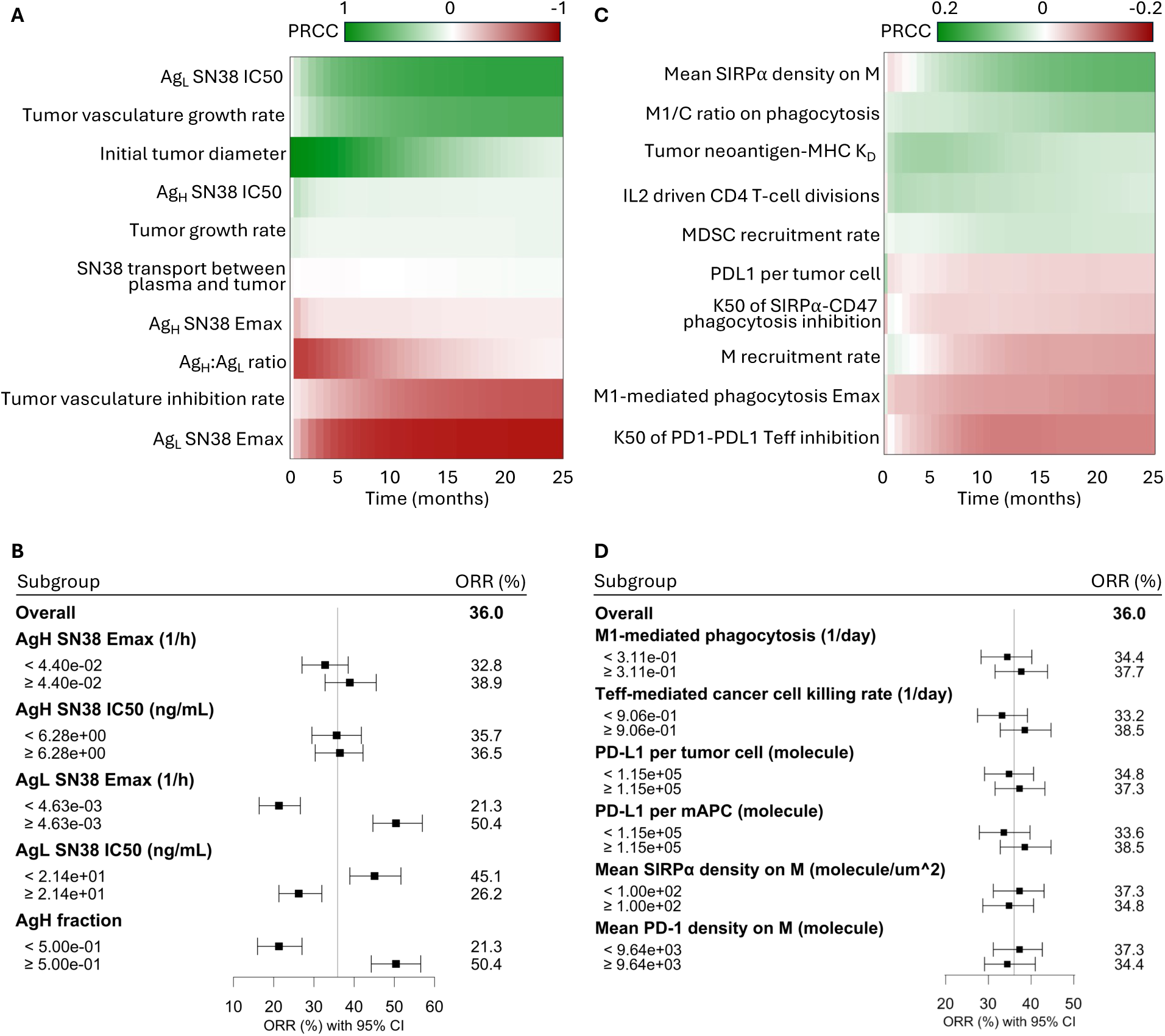
Sensitivity analysis and subgroup stratification of model parameters governing treatment response. (A) Partial rank correlation coefficients (PRCCs) between tumor microenvironment, antibody–drug conjugate (ADC), and free SN-38 pharmacokinetic/pharmacodynamic parameters and model-predicted tumor volume over time. Positive and negative PRCC values (green and red, respectively) denote parameters positively or negatively correlated with final tumor volume, indicating that increases in positively correlated parameters are associated with greater tumor burden, whereas increases in negatively correlated parameters are associated with tumor reduction. (B) Forest plot of model-predicted overall response rate (ORR, %) with 95% confidence intervals (CIs) for the entire virtual patient cohort and for subgroups stratified by parameter-specific median values. The solid vertical line indicates the overall predicted ORR of 36.0%. (C) PRCCs between immune-related parameters and model-predicted tumor volume over time. PRCC color scale was adjusted compared to (A) to improve visibility. (D) Forest plot showing the corresponding model-predicted ORR (%) with 95% CIs for subgroups stratified by median values of immune-related parameters.

Subsequent analysis evaluated the impact of parameter variability on ORR with the population stratified by median parameter values (Fig. 5B). The analysis revealed that efficacy was strongly driven by intrinsic payload sensitivity: ORR was 45.1% versus 26.2% for virtual patients with high versus low Ag_L_ SN-38 potency, respectively. Similarly, high Ag_L_ SN-38 E_max_ was associated with a higher ORR (50.4% versus 21.3%). Furthermore, high Ag_H_ fraction (≥0.5) subgroups demonstrating improved ORR of 50.4% compared to its lower counterpart (21.3%).

These trends are further illustrated at the individual virtual patient level in the waterfall plots (Fig. 6). Virtual patients with low Ag_L_ SN-38 IC50 and high Ag_L_ E_max_ showed greater tumor shrinkage; whereas, stratification by Ag_H_ SN-38 IC50, Ag_H_ E_max_, or TROP2 expression resulted in more evenly distributed tumor responses around the median. In contrast, virtual patients with Ag_H_ fraction values above the median (>50%) exhibited a higher degree of tumor shrinkage.

**Fig. 6.**
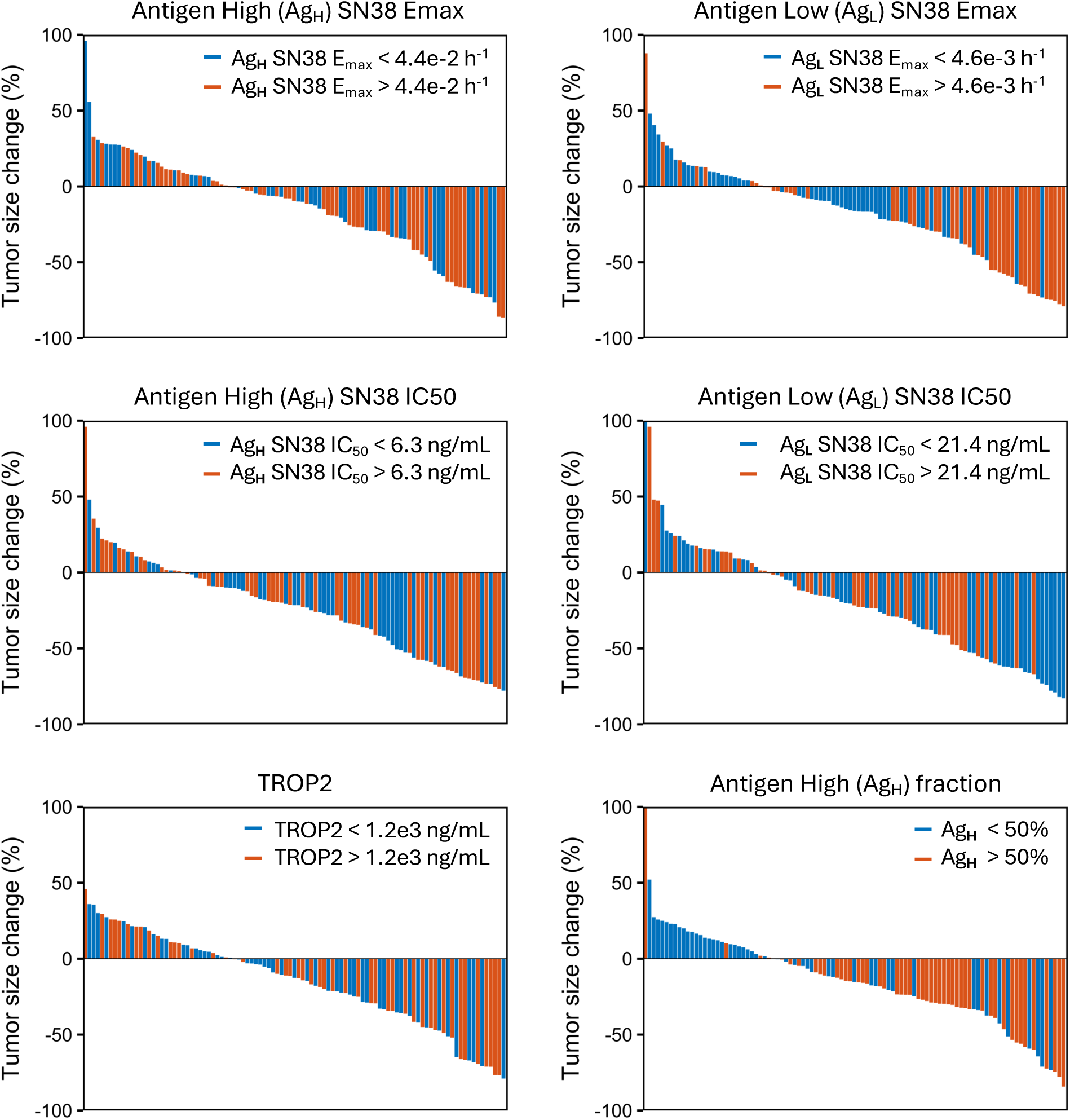
Best overall response in model-predicted tumor diameter for 100 randomly selected virtual patients, stratified by parameter-specific median values.

#### 3.3.2. Sensitivity Analysis of Tumor-Immune Interactions

The sensitivity of the SG therapy on tumor-immune interaction was assessed via evaluating the contribution of macrophage dynamics and checkpoint engagement (Fig. 5C). In this analysis, the maximal macrophage phagocytosis rate emerged as the predominant positive regulator of efficacy with PD-L1 expression per tumor cell also exhibiting a notable positive association. In contrast, parameters associated with immune inhibition, specifically mean SIRPα density on macrophages, demonstrated positive correlations with tumor volume.

Stratification of the cohort by immune parameters further elucidated their impact on clinical response (Fig. 5D). Variability in macrophage activity and checkpoint expression influenced outcomes within the evaluable population; however, these effects were modest and did not substantially alter overall response. Virtual patients characterized by elevated phagocytosis rates (≥ 3.11 × 10⁻¹ day⁻¹) achieved a slightly higher ORR of 37.7%. High baseline PD-L1 expression per tumor cell (≥ 1.15 × 10⁵ molecules) was associated with an ORR of 37.3% versus 34.8% in the low baseline group. Conversely, increased PD-1 density on macrophages (≥ 9.64 × 10³ molecules) yielded an ORR of 34.4% compared with 37.3% in the low-density group, consistent with its inhibitory role on macrophage function, though the overall impact remained limited.

### 3.4. Tumor Microenvironment Immune Activity and Signaling Molecule Analysis

Tumor microenvironment distributions of immune cell populations and signaling molecules were compared between responder and non-responder virtual patients (Fig. 7). Responders exhibited higher densities of effector T cells (T_eff_), regulatory T cells (T_regs_), and helper T (T_h_) cells compared with non-responders. Although T_regs_ and T_h_ cells are often associated with immunosuppressive functions, they do not play a dominant role in driving therapeutic efficacy in this context as SG administration is the primary driver of tumor cell killing. The elevated levels of these T cell populations in responders arise from increased cancer cell death leading to enhanced antigen release. Antigens are taken up by antigen-presenting cells, resulting in increased T cell activation in the lymph node compartment, and subsequent migration of activated T cells to the tumor via the central compartment. This mechanism is supported by simulations in the absence of SG therapy where reduced tumor cell killing leads to lower antigen release and correspondingly lower T cell levels.

**Fig. 7.**
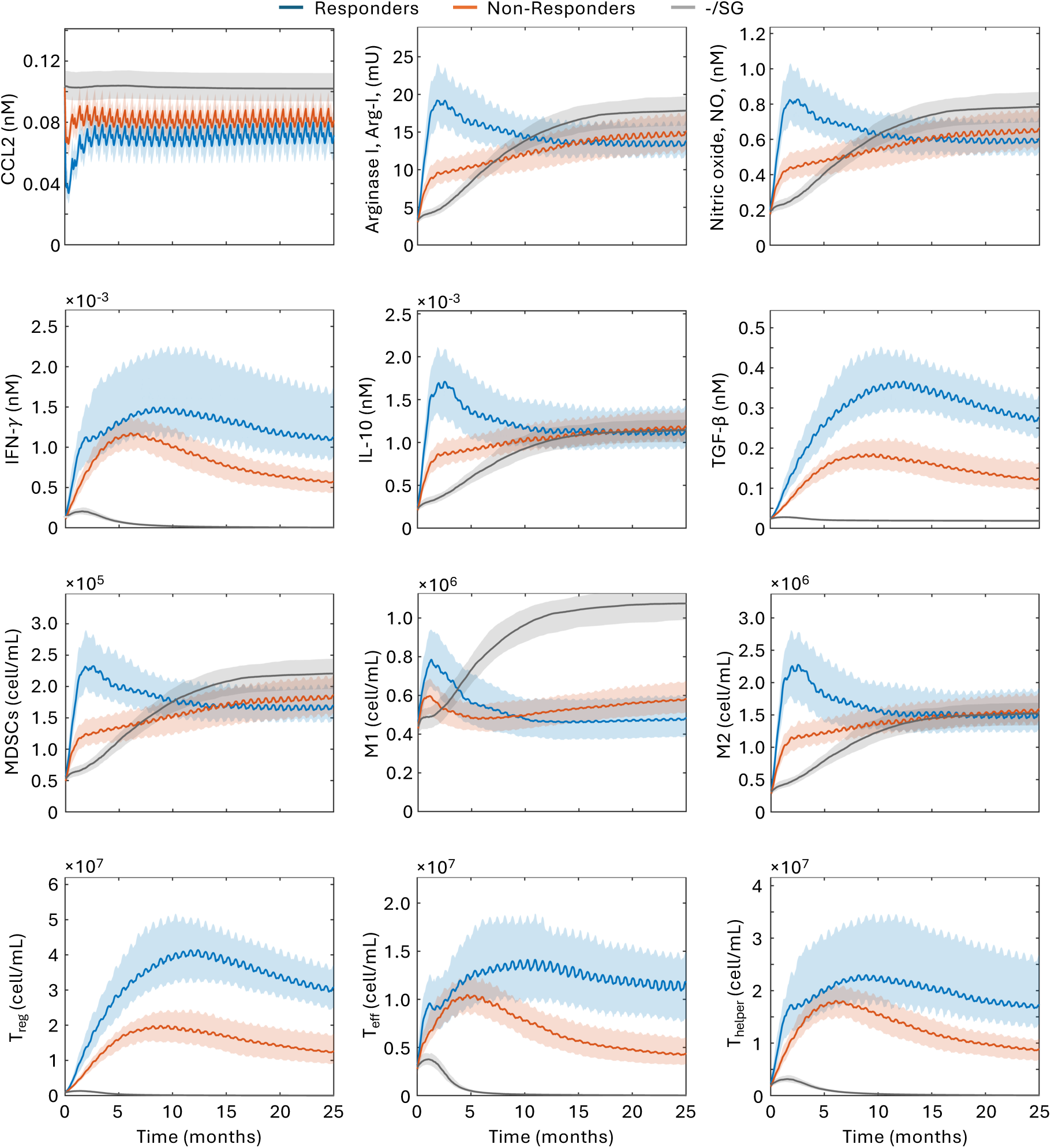
Tumor microenvironment immune activity and signaling molecule analysis in responders and non-responders to SG therapy. Distributions of tumor microenvironment immune activity and signaling molecules across virtual patients categorized as responders (blue) and non-responders (orange) over the course of SG therapy (25 months). Values in the absence of therapy are shown as a reference (gray). Shaded regions represent bootstrap confidence intervals calculated from 244 virtual patients.

The time-dependent IFN-γ profile closely follows that of T_h_ cells consistent with IFN-γ secretion by T_h_ cells in the model (Fig. 7). CCL2 dynamics depend on the number of cancer cells; therefore, SG-treated responders and non-responders both exhibit a rapid decrease in CCL2 levels relative to the no-treatment condition due to reduced tumor burden. Myeloid-derived suppressor cell (MDSC) numbers are driven by CCL2 concentration in the model. While total MDSC numbers are lower in responders (not shown), MDSC density is higher due to early tumor volume shrinkage during therapy. Arginase I (Arg-I) and nitric oxide (NO) follow similar trends as they are secreted by MDSCs. M1 macrophage dynamics also depends on CCL2 concentration resulting in a profile similar to MDSCs. Polarization from M1 to M2 macrophages is driven by M1 abundance as well as TGF-β and IL-10 concentrations. Higher TGF-β levels in responders, secreted by T_regs_, together with elevated IL-10 levels secreted by M2 macrophages lead to increased M2 macrophage densities compared with non-responders and the no-treatment group. Collectively, these results demonstrate distinct tumor microenvironment profiles between responder and non-responder virtual patients throughout the SG therapy.

## 4. Discussion

Antibody-drug conjugates have emerged as promising therapeutics in oncology; however, their clinical outcomes exhibit significant variability among patients. This variability highlights the need for platforms that can mechanistically explore ADC efficacy and optimize their design and administration protocols. Moreover, current efforts to combine ADCs with checkpoint inhibitors necessitate modeling frameworks capable of capturing both ADC pharmacology and tumor-immune interactions simultaneously. In this study, we developed an ADC module and integrated it into our previously established QSP immune-oncology model for TNBC enabling investigation of ADC mechanism of action and the interaction between ADCs and tumor-immune dynamics. While we focused on TROP2-targeted sacituzumab govitecan, the model framework is adaptable to other targets and other ADCs.

TNBC is a heterogeneous tumor with well-characterized molecular subtypes exhibiting distinct antigen expression and payload sensitivity. Antigen expression influences ADC distribution within the tumor microenvironment and payload localization; whereas, cell-intrinsic payload sensitivity impacts cytotoxicity and eventually overall efficacy. Our model accounts for both aspects by representing two distinguished TNBC molecular subtypes using two cell clones that was informed by prior studies characterizing TROP2 expression and SN-38 sensitivity in TNBC.

The mechanism of action of SG has been debated with some studies describing it as more akin to a prodrug than a conventional ADC due to its relatively unstable linker.^45,46^ The half-life of SG (∼16 hours) is considerably shorter than that of other FDA-approved ADCs. TROP2 surface receptor density is approximately ten-fold lower than that of HER2 in cell lines from HER2 amplified patient. Additionally, SN-38 is less potent than typical ADC payloads. Collectively, these SG characteristics imply that its therapeutic efficacy requires SG delivery at higher rates, over shorter durations, and at higher concentrations compared to other ADCs. Nevertheless, the combination of a higher administered dose (10 mg/kg which is 2–4 times higher than other ADCs) and a high drug-to-antibody ratio (DAR = 7.6) is sufficient to achieve meaningful clinical efficacy in TNBC patients. Furthermore, another important factor is the plasma SN-38 levels which in population pharmacokinetics experiments reaches concentrations (∼100 ng/mL) that are 10-100 fold higher than other ADC payload plasma concentrations and above SN-38’s in vitro IC50 values (3–50 ng/mL) suggesting plasma SN-38 may contribute to overall efficacy especially in TROP2-low tumor regions. In vivo studies have shown that SN-38 can be detected in regions where intact ADC is absent.^47^ This could happen either as a result of plasma dissociation or intracellular cleavage followed by efflux to neighboring cells, potentially contributing to efficacy via a bystander effect. However, previous theoretical analyses suggest that the high diffusivity of SN-38 within the tumor microenvironment would lead to substantial plasma washout thereby limiting the contribution of the bystander effect.^48^

Our model explicitly accounts for the impact of intratumoral heterogeneity on ADC efficacy. When Ag_H_ cells represent a minority of the tumor population, they are depleted more rapidly, amplifying the overall loss of cytotoxic effect. Fig. 3 illustrates this heterogeneity by showing how the Ag_H_:Ag_L_ ratio influences therapeutic response. This effect may arise either from differences in targeted payload delivery via SG-TROP2 binding or from variation in the fraction of payload-sensitive versus resistant tumor cell clones. Targeted SG delivery contributes only a small fraction of total intracellular SN-38 exposure (Fig. 2), suggesting that differences in cell-intrinsic sensitivity are the dominant mechanism. In this scenario, plasma-derived unconjugated SN-38 diffuses into the tumor microenvironment acting on both Ag_H_ and Ag_L_ cells and serving as the primary driver of efficacy. Consistent with this scenario, prior studies have shown inhibiting epithelial-to-mesenchymal transition (EMT) factors enhances SG efficacy not only by increasing TROP2 expression and ADC uptake but also by sensitizing tumor cells to SN-38.^20^ Together, these results underscore the importance of considering heterogeneity in both antigen expression and payload sensitivity. In SG treated TNBC, these factors are correlated making mechanistic modeling essential for disentangling their individual contributions and quantifying their effects on ADC therapeutic efficacy.

SG payload SN-38 has been utilized in clinical practice as its prodrug irinotecan which is enzymatically converted from an inactive precursor to SN-38. Irinotecan has demonstrated clinical efficacy across several solid tumors; however, its use is limited by dose-related toxicities such as neutropenia and diarrhea which correlate with SN-38 systemic exposure (e.g., ∼229– 474 ng·h/mL over 0–24 h depending on dose).^49–51^ To improve irinotecan’s pharmacokinetics and potentially reduce adverse events, other prodrugs have been developed such as EZN-2208 (PEGylated SN-38) to prolong circulating SN-38 exposure.^52,53^ Approximately 9 mg/m² EZN-2208 was administered once every three weeks and showed an ORR of 22.5 % in previously treated TNBC patients (11 of 49). The administered EZN-2208 dose was considerably lower than SG (10 mg/kg on days 1 and 8 of a 21-day cycle) and as a result EZN-2208’s SN-38 exposure levels were 4–5 times lower than those observed with SG (AUC_SG_: ∼3,696 ng·h/mL vs AUC_EZN2208_: ∼1,000 ng·h/mL). In parallel with this lower exposure, the efficacy of EZN-2208 in early clinical trials was somewhat lower than SG (ORR: 22.5 % vs 31.4 %). Nevertheless, these results demonstrate that SN-38 prodrugs can achieve considerable antitumor activity. Collectively, these findings support the concept that administering SN-38 as a prodrug can be an effective therapeutic strategy and highlight that the prodrug characteristics of SG likely contribute to its efficacy in TNBC.

Antibody–drug conjugate (ADC) and immune checkpoint inhibitor combination therapies have received increasing attention in recent years, with several clinical trials currently ongoing. Notably, results from the SG–pembrolizumab combination trial have recently been reported.^54^ Motivated by these findings, we analyzed the effects of SG therapy on immune activity within the tumor microenvironment. Our simulations demonstrated increased T cell concentrations in the tumor microenvironment following SG treatment compared with no therapy in both responder and non-responder virtual patients. This increase is driven by enhanced therapy-mediated cancer cell killing, resulting in greater antigen release and subsequent immune activation. These insights are particularly relevant considering the SG–pembrolizumab clinical trial results, which showed improved objective response rate (ORR) and progression-free survival (PFS) compared with SG or pembrolizumab monotherapy. Our results (Fig. 7) suggest that these efficacy gains may arise from mechanistic synergy between the two agents, whereby SG-induced tumor cell killing increases effector T_eff_ activity in the tumor microenvironment and pembrolizumab-mediated checkpoint inhibition further amplifies T_eff_-mediated tumor cell killing. Further investigation is required to quantitatively evaluate this synergy and to formally assess combination therapy effects within the model framework.

Several limitations should be considered when interpreting the results of this study. First, the effect of the payload on immune cells was not incorporated due to limited available evidence. Second, the glucuronidation of SN-38 within cancer cells was previously reported, which could affect local drug availability, but was not accounted for due to lack of quantitative data. Third, the development of resistance over time was not included, and the growth rates of different tumor cell clones were assumed to be identical due to lack of detailed information. Fourth, the model did not incorporate the formation of new lesions arising from metastasis. Fifth, free antibody binding was assumed to be negligible, as most of the ADC was reported to remain conjugated throughout the experimental analyses. Additionally, the initial calibration was performed using Phase I/II basket trial data and validated against Phase III clinical trial results. For SG, these trials yielded similar outcomes despite originating from different lines of therapy. While this approach provided a reasonable estimation of efficacy when extrapolating from early-phase to late-phase trials, further work is needed to mechanistically account for differences in lines of therapy. Lastly, some studies suggest that TROP2 blockade can inhibit tumor growth through signaling pathways affecting proliferation and survival, highlighting the role of TROP2 as an oncogene. The impact of this mechanism on SG efficacy was not addressed due to the lack of relevant experimental data.

## Study Highlights

### What is the current knowledge on the topic?

TROP2-targeted antibody–drug conjugates (ADCs) have demonstrated clinical efficacy in multiple solid tumors and emerging clinical data suggest improved outcomes when combined with immune checkpoint inhibitors. The mechanistic drivers of response heterogeneity and immune modulation by ADCs remain poorly understood.

### What question did this study address?

This study developed a mechanistic TROP2-targeted ADC model integrated into a quantitative systems pharmacology (QSP) immuno-oncology framework for triple-negative breast cancer (TNBC). Using sacituzumab govitecan (SG) as an application example, determinants of response and tumor microenvironment immune dynamics were investigated while explicitly accounting for TNBC heterogeneity in TROP2 expression and intrinsic payload sensitivity.

### What does this study add to our knowledge?

The model identifies plasma payload levels and intrinsic payload sensitivity as dominant drivers of efficacy and demonstrates that SG-mediated tumor cell killing enhances antigen-driven T cell activity in the tumor microenvironment.

### How might this change drug discovery, development, and/or therapeutics?

These insights provide a mechanistic basis for optimizing ADC-checkpoint inhibitor combination strategies and a framework to quantitatively evaluate combination therapies.

## Availability of data and material

All data generated or analyzed during this study as well as the model in SBML format are included in this published article and its supplementary information files.

## Authors’ contributions

Study conception and design, all authors; Model implementation and testing, SD; Data collection and analysis, SD, ASP; Writing Original Draft, SD; Revising and editing manuscript, all authors. All authors read and approved the final manuscript.

## Acknowledgements

We thank Dr. Ravi Varadhan for his expert guidance on the statistical methodology used to determine the required virtual patient sample size for PRCC analysis.

## AI Use Disclosure

Language editing assistance was provided using an artificial intelligence-based writing tool (ChatGPT, OpenAI, GPT-5) to perform grammatical corrections and improve readability. The AI system was prompted only to check grammar and improve readability, without altering or contributing any technical or scientific content, data interpretation, or conclusions. No AI tools were used for figure generation, content creation, or any other aspect of manuscript preparation. Potential biases and limitations of AI use were negligible, as edits were restricted exclusively to language refinement.

