## Supplementary material for "Quantitative Systems Pharmacology Model for TROP-2 Targeted Antibody-Drug Conjugate in Triple-Negative Breast Cancer": Supplemental_Mateial_Quantitative Systems Pharmacology Model for TROP-2 Targeted Antibody-Drug Conjugate in Triple-Negative Breast Cancer.docx

Aleksander S. Popel^1,3^

^1^Department of Biomedical Engineering, School of Medicine, Johns Hopkins University;

^2^Merck & Co., Inc., Rahway, NJ, USA;

^3^Sidney Kimmel Comprehensive Cancer Center, Johns Hopkins University

*Corresponding author:

Sedat Dogru, Ph.D.

Department of Biomedical Engineering

School of Medicine

Johns Hopkins University

Baltimore, MD 21205

**1. ADC Module**

**1.1. Parameterization of ADC-TROP2 and payload kinetics**

In the ADC module, both the antibody-drug conjugate (ADC) or free payload can be delivered from the central compartment to the tumor compartment. Within the tumor, the ADC binds to its target antigen, TROP2. The ADC-TROP2 complex undergoes internalization, followed by intracellular cleavage and release of the cytotoxic payload (Fig. S1). After cleavage, both the antibody and receptor are assumed to be degraded. Intracellular payload, delivered either through ADC-TROP2 internalization or via influx of free payload from the tumor microenvironment induces cytotoxicity. Cytotoxic effects are modeled using a Hill function characterized by a maximum cytotoxicity rate (E_max_) and a half-maximal inhibitory concentration (IC₅₀). These parameters are defined separately for the two tumor cell clones.

The equilibrium dissociation constant (K_D_) for TROP2-SG binding was obtained from literature.^1^ Association (k_on_) and dissociation (k_off_) rate constants were determined by modeling TROP2-SG binding and internalization dynamics using published surface binding datasets that reported both cell-surface and internalized SG levels (Fig. S2).^2^ The internalization rate (k_int_) and cleavage rate (k_cleave_) were also obtained from literature measurements.^2^ Following SN-38 release, both TROP2 and the antibody are assumed to undergo intracellular degradation. TROP2 surface expression is assumed to remain constant in the absence of SG as implemented in similar modeling studies.^3,4^ SN-38 influx and efflux rate constants were obtained from published experimental data.^5,6^

Payload, SN38, IC₅₀ values were obtained from The Genomics of Drug Sensitivity in Cancer Project (GDSC) database for TNBC cell lines, and the median value was calculated for each molecular subtype.^7^ E_max_ values were determined using in vivo tumor growth inhibition data reported in the literature for two representative cell lines corresponding to antigen-high and antigen-low phenotypes: MDA-MB-468 and MDA-MB-231, respectively.^8,9^ First, a pharmacokinetic (PK) model was developed to reproduce the pharmacokinetics of sacituzumab govitecan (SG) in mice serum and tumor (Fig. S3A-B). The saline (no-therapy) control group was then used to characterize baseline tumor growth dynamics (Fig. S3C-D). Subsequently, model calibration was performed using the SG treatment group to estimate the maximum cytotoxicity rate (E_max_) for both cell types, while employing IC₅₀ values obtained from the database for each cell line.^7^ The estimated E_max_ values were directly translated to the human simulations without additional scaling, as cytotoxicity was assumed to be cell line–specific rather than species-dependent.

**1.2. TNBC heterogeneity:** TROP2 expression levels for the cell lines, categorized as basal-like or mesenchymal-like subtypes, were obtained from the Human Protein Atlas database (Fig. S4).^10^ Our analysis indicated that basal-like cell lines consistently exhibit higher TACSTD2 expression. Gene expression values were then approximated to cell-surface TROP2 levels using previously reported membrane protein quantifications in selected cell lines.^11^

**2. Determination of Virtual Patient Sample Size for Calibration and Sensitivity Analysis**

The effect of the number of virtual patients (VPs) on response status was evaluated. Results indicated that beyond 400 VPs, no significant differences in response status were observed (Fig. S5A). Therefore, model calibration was performed using 400 virtual patients.

Partial rank correlation coefficient (PRCC) analysis was conducted to quantify parameter sensitivity. To determine the VP population size required for stable PRCC results, a top-down concordance coefficient (TDCC) analysis was performed across increasing VP sample sizes.^12^ TDCC analysis demonstrated that PRCC-based sensitivity rankings stabilize for VP populations greater than 1200 (Fig. S5B). Accordingly, all subsequent sensitivity analyses were performed using 1200 virtual patients.

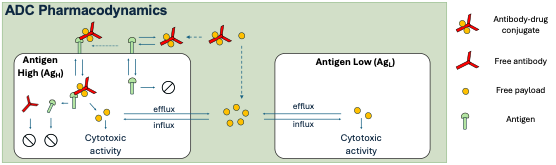

**Fig. S1** Schematic of the ADC module. The diagram illustrates key processes including ADC–TROP2 binding, internalization of the ADC–TROP2 complex, intracellular payload release via cleavage, cytotoxic action of the payload, and efflux/influx of free payload in the tumor microenvironment.

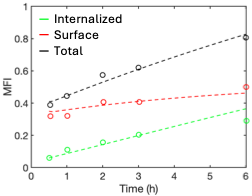

**Fig. S2** Estimation of TROP2-SG binding kinetics parameters using experimental data. Scatter points represent cell-surface, internalized and total SG–TROP2 complex mean florescence intensity (MFI) levels measured experimentally.^2^ Model curves were fitted to these data to determine the TROP2-SG association and dissociation rate constants.

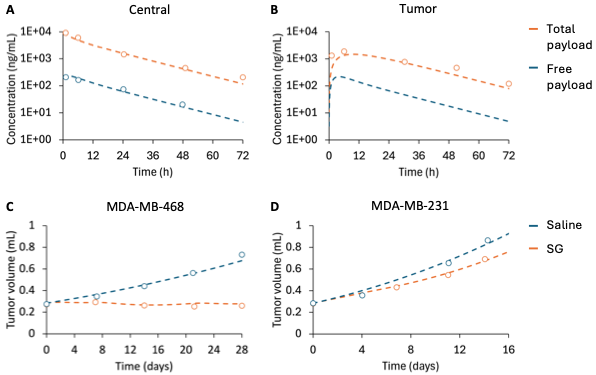

**Fig. S3** Preclinical modeling of SG therapy for parameter estimation. The pharmacokinetic (PK) module was optimized to reproduce preclinical measurements of (A) central and (B) tumor concentrations of total and free payload. Note that free payload concentration in tumor has not been reported in the literature. Using the optimized PK module, tumor growth inhibition was simulated for (C) MDA-MB-468 and (D) MDA-MB-231 cell lines, representing the sensitive (antigen-high, Ag_H_) and resistant (antigen-low, Ag_L_) clones in the model, respectively, to estimate the maximum cytotoxicity rate (E_max_) for each clone.

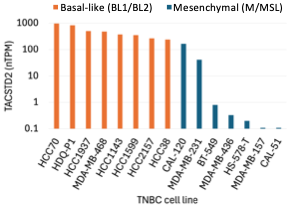

**Fig. S4** TROP2 (TACSTD2) gene expression levels in TNBC cell lines, stratified by molecular subtype. Expression is shown in normalized transcripts per million (nTPM).

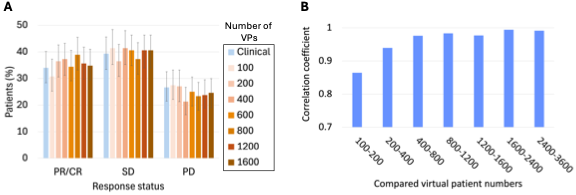

**Fig. S5** Effect of virtual patient (VP) population size on response status and global sensitivity analysis. (A) Response status outcomes stabilized beyond 400 VPs, supporting the use of 400 VPs for model calibration. (B) A top-down concordance coefficient (TDCC) analysis demonstrated stabilization of partial rank correlation coefficient (PRCC) sensitivity rankings for VP populations greater than 1200. Therefore, simulations were performed with 1200 virtual patients.
